# Impact of sequencing technologies on long non-coding RNA computational identification

**DOI:** 10.1101/2022.04.15.488462

**Authors:** Alisson G. Chiquitto, Lucas Otávio L. Silva, Liliane Santana Oliveira, Douglas S. Domingues, Alexandre R. Paschoal

## Abstract

The correct annotation of non-coding RNAs, especially long non-coding RNAs (lncRNAs), is still an important critial challenge in genome analyses. One crucial issue in lncRNA transcript annotation is the transcriptome resource that supports lncRNA loci. Long-read technologies now bring the potential to improve the quality of transcriptome annotation. Consequently, long non-coding RNAs (lncRNA) are probably the most benefited class of transcripts that would have improved annotation using this novel technology. However, there is a gap regarding benchmarking studies that highlighted if the direct use of lncRNA predictors in long-reads makes more precise identification of these transcripts. Considering that these lncRNA tools were not trained with these reads, we want to address: how is the performance of these tools? Are they also able to efficiently identify lncRNAs? We could provide evidence of where and how to make potential better approaches for the lncRNA annotation by understanding these issues. Keywords: Non-coding RNAs, high-throughput sequencing technologies, coding, methods, benchmarking, tools, NGS, transcripts

## Introduction

High-throughput sequencing technologies (HTS) opened up a new era in omics analysis. Short-read-based transcriptome-reconstruction methods delivered large annotations with relatively few resources. Although a myriad of annotation efforts were possible due to HTS, the quality of reconstructed transcripts is often doubtful because of the inherent difficulty of reconstructing transcript structures from shorter sequence reads [11]. This means that long-read sequencing techniques, like Iso-seq and Nanopore, have a great potential to improve the quality of transcriptome annotation.

Even with improvements in annotation, most efforts are aimed to better investigate gene-coding transcripts. Long non-coding RNAs (lncRNAs) still lack behind a dedicated annotation effort [2, 13, 15]. In this sense, lncRNA identification is probably the most benefited class of transcripts that would have improved annotation using long read sequencing technologies. The GENCODE project [7], as a foundational resource for human genome annotation, has recently recognized the benefits of using long-read transcriptome sequencing [7]. In the human transcriptome, the benefits of annotating lncRNAs using long-read transcript data have already been proved as an important advance [7].

The most recent release of lncRNA annotation of GENCODE is now heavily based in long-read sequencing analysis, which makes this database as a reference resource to evaluate the performance of lncRNA computational predictors. Most of them were developed to be applied in reconstructed transcripts from short reads. In this sense, we were prompted to test if lncRNA predictors are efficient to correctly identify lncRNAs in actual data that were developed in long-read technologies, namely ISO-seq (PacBio) and Nanopore, with an emphasis in plants, in which similar efforts were not addressed.

Our objective was also to transfer this knowledge to two other real-world cases: can lncRNA identifiers be directly used in long-reads to make a more direct identification of these transcripts? Are they also able to efficiently identify plant lncRNAs, where most computational approaches are not well trained?

Understanding these issues, we could provide evidence of where and how to make potential better approaches for the lncRNA annotation.

## Materials and Methods

We have conducted an review of the lncRNAs classification methods, in which several approaches have been developed. For better understanding, Supplementary Table S1 presents the main characteristics of these works.

In 6, the authors selected several popular or newly published software packages covering most of the statistical models and analysis algorithms commonly used in lncRNA recognition. From this work, we selected CNCI [19], COME [8], CPAT [21], CPC2 [10], lncScore [25], PLEK [12], PLncPro [18] and RNAplonc [14]. The other tools mentioned in 6 had problems during execution and we did not list them in this work. Finally, LncADeep [24], LncRNAnet [1], CREMA [17], LncMachine [3], RNAmining [16] and RNAsamba [4] are referenced in related works.

In general, the analyzed tools use supervised methods of machine learning using binary classification (lncRNAs and protein-coding genes (mRNA)). Regarding the approach used for feature extraction (Approach column), we observe a large domain of approaches based on biological features, such as ORF and sequence structure descriptors. On the other hand, few works have explored mathematical approaches to feature extraction, such as complex networks. Finally, hybrid approaches bring characteristics of both mathematical and biological approaches.

All the tools used receive a FASTA format sequence as input. Only the COME tool [8] needs the GTF file, which we extracted from the GENCODE website.

### Datasets used for analysis

A total of 25 datasets were used, of which 17 of plants and 8 were of *Homo sapiens* (Supplementary Tables S4). We opted to restrict our plant analyses to angiosperms with represerntative long read transcriptome data. One monocotyledonous species (*Triticum aestivum*), one dicotyledonous species (*Arabidopsis thaliana*), and one basal species (*Amborella trichopoda*) with data in more than one database were chosen.

#### Plant transcript and lncRNA data

We used short-read data from CANTATAdb [20] and NONCODE [26] non-coding RNAs databases. In particular, we used lncRNAs of *Amborella trichopoda* and *Arabidopsis thaliana* from CANTATAdb 2.0 [20], as a curated set of previously annotated lncRNAs. Since *Triticum aestivum* is not represented in CANTATAdb, we used data from NONCODE [26], a database of non-coding RNAs based on short reads. Next, we used two datasets to represent a real-world case of long-read data to employ in a direct analysis of lncRNA predictors:

1. ISO-Seq data were obtained using the ISOdb database (obtained in [23] with de respective IDs). We obtained two datasets for*Amborella trichopoda* (IDs: SAMN06320627 and SAMN06320628), six for *Arabidopsis thaliana* (IDs: SAMN04456597, SAMN04456598, SAMN04456599, SAMN04456600, SAMN04456601 and SAMN04456602), and three for *Triticum aestivum* (IDs: SAMN04456603, SAMN04456604 and SAMN04456605). Since these species represent a good diversity in Angiosperms, they were our focus in previously annotated lncRNAs.
2. The *Arabidopsis thaliana* is the only plant species with public deposited transcriptome datasets generated by NANOPORE [5] where the data were extracted from three samples in NCBI Sequence Read Archive (SRA) (GSE141641 - IDs: SRR10611193, SRR10611194 and SRR10611195 by using fastq-dump tool) (Supplementary Tables S4).

### Homo sapiens transcript and lncRNA data

In GENCODE [7], we used datasets of long non-coding RNAs from a version based on short reads (version 21, October 2014) and one based on long reads (version 38, May 2021). We used two datasets to represent a real-world case of long-read data to employ in a direct analysis of lncRNA predictors:

1. A dataset which contains a direct sequencing of RNA and cDNA of a human transcriptome on the MinION and GridION platforms [9], was used as a reference benchmark for sequencers on the Nanopore platform. We used this dataset to assess a real-world case of predictor application (available in [22] - NA12878 - NANOPORE file NA12878-DirectRNA.pass.dedup.fastq.gz)
2. Iso-Seq reads were also retrieved from ISOdb database into the Donwload section [23], where we obtained three *Homo sapiens* datasets (IDs: SAMN00001694, SAMN00001695, and SAMN00001696).

### Pre-processing, parse and data analysis

Figure S1 summarizes all of the processes carried out during this study.

We needed to manually remove a single sequence from the GENCODE dataset (ID: ENST00000625083.1) because it included N bases. All lncRNA prediction tools were run on lncRNAs from versions 21 and 38 of GENCODE. Based on average sensitivity (Figure S5), runtime [6], and the size of the dataset, we selected tools for running on long-read transcritptome data. We ran four tools in Iso-seq data (CNCI, LncADeep, lncRNAnet, and PLEK), and two in Nanopore data (LncADeep and PLEK).

It is important to highlight that in plants, PLncPro tool has two models (dico and mono). In this case, the mono model was used for *Triticum aestivum*, the dico model for *Arabidopsis thaliana*, and both models were used for *Amborella trichopoda*.

To evaluate the tool’s performance on datasets that contain only lncRNA sequences(lncRNA catalogs - lncRNAs in Data Type column - in Supplementary Tables S2), we used the Sensitivity Equation, the same one used to calculate the performance of predictive models. For the calculation, we considered as true positives (TP) the lncRNA sequences that were classified as lncRNA, and for lncRNA sequences classified as mRNA we considered them as false negatives (FN). Finally, by applying these values to Equation 1, we were able to obtain the tool’s sensitivity value. We get the number of zero False Positives (FP) and zero True Negatives (TN) on datasets that contain only lncRNA sequences, then specificity and other commonly used values in this situation were not calculated as we do not have mRNA sequences in these datasets.

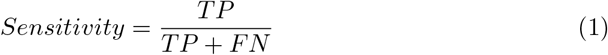

Relative frequency is a measure we use to calculate the proportion of lncRNA over a set of RNA sequences. We obtain the amount of lncRNA given by the tools and divide it by the total number of sequences in the dataset, according to Equation 2. The relative frequency was calculated on the datasets that contained RNA data (Transcriptome in Data Type column - in Supplementary Tables S2).

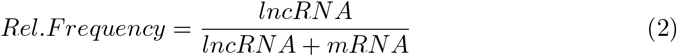

## Results

We run our pipeline and tools (Figure S1) to calculate Sensitivity for the specialized lncRNA databases (GENCODE, CANTATAdb, and NONCODE).

### Plant data

The data obtained for plants were analyzed using CPC2, Crema, lncMachine, PLncPro and RNAplonc tools (Figures S2, S3 and S4). In the case of short reads (Figure S2), the PLncPro monocot (PLncPro-mono) and dicot (PLncPro-dico) models were used for *Triticum aestivum* and *Arabidopsis thaliana*, respectively, while both models were used for *Amborella*. Except for CREMA, all tools attained results of over 90% and were close to the sensitivity values in terms of the decimal places. It is important to note that CPC2 was the best tool in two of the three tests, but it only determines whether a sequence is coding or non-coding, meaning that an additional tool is needed to analyze the type of non-coding sequence. Of the tools, the CPC2 was the best in *T. aestivum* and *Amborella*, and lncMachine was the top performer for *A. thaliana*. Finally, both CPC2 and RNAplonc exhibited a high degree of stability (*>*96%) even when the plant was changed. In other words, these tools can be used for any angiosperm taxon.

The 11 plant datasets from ISOdb were analyzed using the five short-read data tools (Figure S3). CREMA continued to have the worst results of all of the programs compared. Of the 11 datasets analyzed, RNAplonc identified the largest number of lncRNAs in six. For *Amborella*, the PLncPro-mono model identified more lncRNAs than the PLncPro-dico model.

We ran the CPC2, lncMachine, and RNAPlonc tools for the Nanopore datasets (Figure S4). The reason for using these tools was that they exhibited a good runtime and one of the best sensitivity values during the execution of the tests with short reads and ISOdb. The number of lncRNA identified by RNAplonc was similar to the CPC2, while lncMachine identified the smallest amount.

### *Homo sapiens* data

Figure S5 provides an overview of the results for the *Homo sapiens* lncRNA data from GENCODE versions 21 and 38, which were compared due to their differences in the use of HTS data.

The two best sensitivity values for v21 were obtained by RNAmining, followed by PLEK. Meanwhile, the best sensitivity values for v38 were achieved by RNAmining and LncADeep. By calculating the average value of the sensitivity metric between the two versions of GENCODE, we found that the five tools with the best average values are RNAmining, CNCI, LncADeep, lncRNAnet, and PLEK.

Some tools have filtering mechanisms in place when entering sequences. This resulted in a smaller number of sequences being analyzed. For example, LncScore only considered sequences longer than 200 nucleotides, meaning that it only processed 26,124 of the 26,414 lncRNA sequences contained in GENCODE version 21.

For ISOdb dataset, the four tools with the best average sensitivity values were run, except for RNAmining due to the fact that we were using RNAmining Web, which has limitations in terms of the size of files. The proportion of lncRNAs identified in the datasets obtained from ISOdb are listed in Figure S6.

For the human Nanopore dataset, we used the tools LncADeep and PLEK (Figure S7). Since the Hsa reference sample dataset contained over 10 million sequences, the tools employed had lower runtimes with good results for datasets obtained from GENCODE. Figure S7 lists the results of the lncRNA identification.

When we observe PLEK tool results in GENCODE v21 dataset (short-reads) in comparison with GENCODE dataset v38 (long-reads), we observe a lower sensitivity value (from 98.28% to v21 to 93.70% to v38 - Figure S5). Also, PLEK had the greater reduction against all tools for lncRNA identification in GENCODE dataset version 38. Finally, PLEK tool was the one that show the lowest relative frequency of lncRNA identification for the real-case long-reads (ISOdb and NANOPORE dataset) - Figures S6 and S7).

### Comparing annotation in human data

After the results were tabulated, we created Upset Plots in order to analyze the results obtained by the tools more clearly. These diagrams used the six tools (disregarding the results of COME) that obtained the highest average sensitivity values, namely the following: CNCI, CPC2, LncADeep, lncRNAnet, PLEK and RNAmining. The diagrams were built using sequence IDs. The results provided by the COME tool were not displayed because the tool uses the GTF format as input and its IDs do not match the IDs of the FASTA files. Thus, it is not possible to intersect the results of the other tools with those of COME.

The diagrams in Figure S8 were made with the sequences that were not classified as lncRNA in the datasets obtained from GENCODE versions 21 and 38, respectively. In Figure S8a, we can see that six sequences were considered to be coding by all six tools. The IDs of these sequences can be found in Supplementary File 2. Figure S8b shows that a total of two of these sequences were considered to be coding by all six tools. The IDs of these sequences can be found in Supplementary File 2.

The analysis of overlapping sequences among all of the top 5 tools used for the coding sequences from GENCODE showed that one of the two transcripts (ENST00000539086.5) of the v38 analysis was superimposed over a region with two coding genes.

## Discussion

With respect to the GENCODE data, version 21 is an older version in which lncRNAs were mostly predicted using short-read data, while version 38 is the most recent and used long-read transcriptome data to predict lncRNAs. Using this difference we addressed: how did predictors fare with these data?

We can interpret the results in 3 ways: i) Tools that had an increase in the Relative Frequency of lncRNA in the analysis of long-reads: The LncADeep, RNAmining and lncRNAnet tools had the greatest increases in the relative frequency of lncRNA when they analyzed long-read sequences. When looking at the results of these tools in the datasets of GENCODE V21 and V38 had the largest increases in Sensitivity being these increases respectively of 1.1%, 0.68% and 0.86%. When we analyze LncADeep in depth, this pattern continues to occur, when we observe the results obtained in the Homo sapiens Datasets of ISOdb and NANOPORE, it is noted that the tool had one of the highest relative frequency of lncRNA, being the first in the NANOPORE dataset and the third in the ISOdb dataset. The same occurred for lncRNAnet, which was the 2nd tool with the highest relative frequency of lncRNA in the ISOdb dataset.

i. Tools that had a drop in the Relative Frequency of lncNRA in the analysis of long-reads: When we compare the results obtained from PLEK on the GENCODE v21 (short-reads) and v38 (long-reads) datasets, we can see a decrease in Sensitivity. This pattern continues to occur when we analyze the results obtained in the Homo sapiens Datasets from ISOdb and NANOPORE, in which PLEK was the tool that had the lowest relative frequency of lncRNA. In the analysis of NANOPORE, PLEK had a relative frequency of lncRNA of 59.15%, and in the analysis of Datasets of ISOdb it had a mean relative frequency of lncRNA of 44.11%.
ii. Tools that did not indicate change in Relative Frequency of lncRNA in long-read analysis: The CNCI, COME, CPAT, lncScore and RNASamba tools did not indicate major changes in the identification of lncRNA when they analyzed long-read sequences, however it is worth mentioning the CNCI tool that obtained the second best average of Sensitivity (97.47%) in the analysis of GENcode V21 and v38 results and was the tool that most identified lncRNA in the ISOdb Homo sapiens datasets, with an average lncRNA frequency of 62.27%.

For plants, it is clear that CREMA did not achieve a satisfactory result in the analyses performed, even according to its technical documentation. However, CPC2 and RNAplonc are more stable for the diverse selection of plants used and were able to identify the most lncRNAs in long-read data in general. Overall, lncMachine identified 10% fewer lncRNAs in NANOPORE than CPC2 and RNAplonc. Of the 13 ISOdb datasets analyzed, RNAplonc identified the largest number of lncRNAs in nine. These results indicate that the combination of CPC2 and RNAplonc have considerable potential for applications in both short and long-read technologies dueto their consistent results across a wide variety of plant taxa. In addition, CPC2 is a tool that classifies your results only as coding or noncoding, while RNAplonc classifies them as mRNA and lncRNA, further indicating the potential of these tools.

## Conclusions

We present a benchmarking of predictors with human and plant data with the aim of analyzing the impact of short and long-read sequencing on the identification of lncRNAs. The lncRNAs prediction tools were demonstrated to be capable of applications involving long-read data from both humans and plants. However, this does not necessarily apply to all tools, as discussed. In sum, we observed an improvement in annotations of lncRNA in humans, especially when using CPAT, RNAmining, lncRNAnet, and LncADeep. Meanwhile, RNAmining, lncRNAnet, LncADeep, and CNCl obtained the best results irrespective of the dataset or analysis carried out on human material. In plants, on the other hand, RNAplonc and CPC2 exhibited the best capacity for adapting to the datasets, even considering the diversity of plants analyzed. A further contribution of this study was the diagrams illustrating overlapping classifications as coding sequences when the dataset was comprised of human lncRNAs. Whether lncRNAs classified in this way truly are coding or whether this is an error across all predictors remains. Finally, we believe that this study has provided points of comparison between the sequencing technologies and datasets with respect to the problem of identifying lncRNAs.

## Author contributions statement

A.R.P. and D.S.D. conceived and designed the study. A.R.P. and D.S.D. oversaw and coordinated the project. A.G.C. and L.O.L.S. developed, implemented, realized the experiments and analyzed the results. All authors wrote this article. All authors approved the final version of this manuscript.

## Additional information

The authors declare that they have no conflict of interest.

## Supplementary Information

### Additional file 1 — Supp. Mat. Table 1-8

All results from performances in each tool applied all datasets described in this report.

### Additional file 2

The data used to create Figure S8 is available at: https://github.com/alerpaschoal/Benchmarking-HTS-LncRNAs-Tools

---

**Figure S1.**
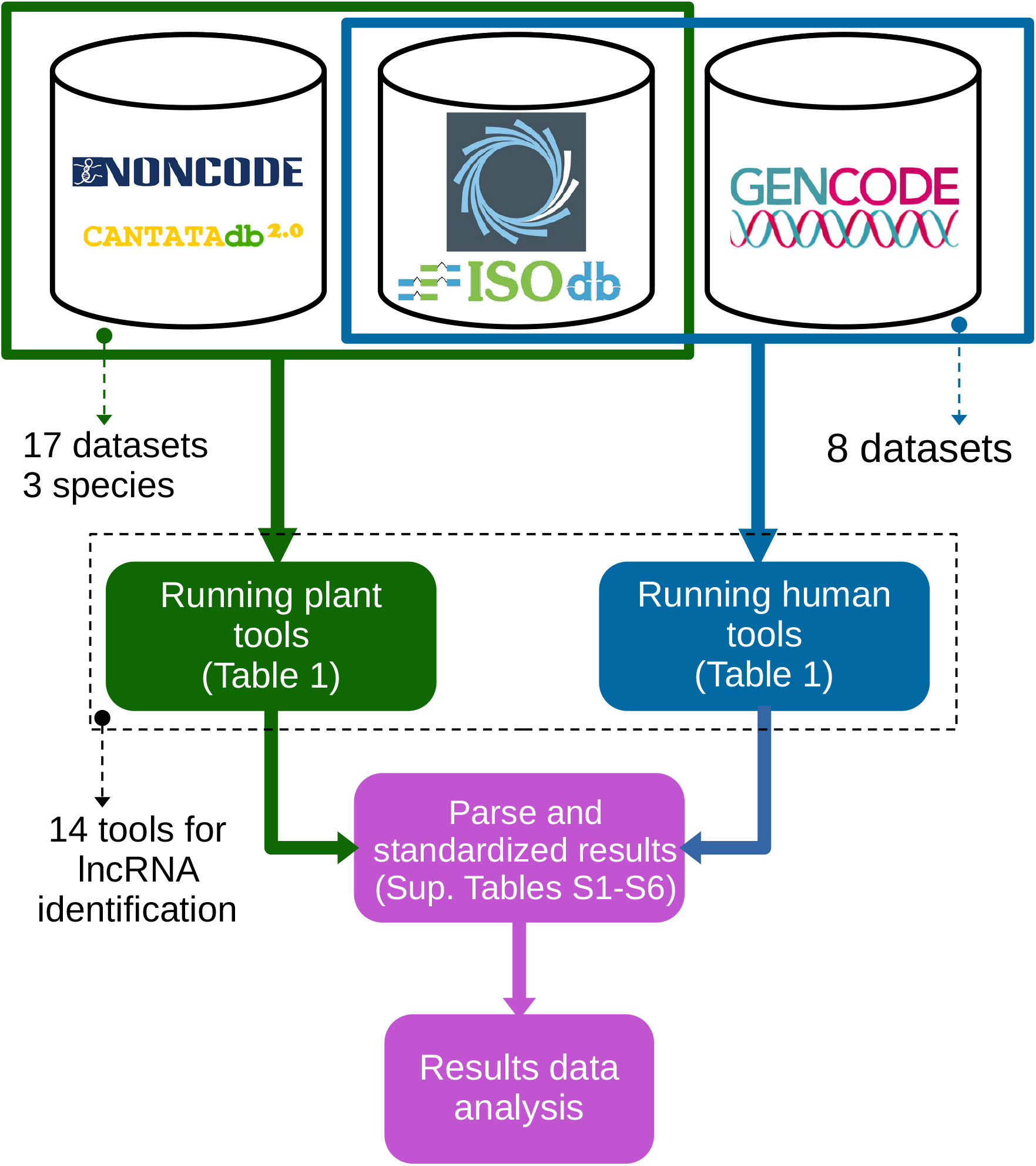
The flowchart of the step-by-step of the analysis done. We obtain the 25 datasets from different sources - 17 for plants of 3 species and 8 for *Homo sapiens*. Then we prepare and run the classification tools on the obtained datasets. Each tool provides the results in a specific format, so we standardize the results (see Supplementary Tables S1 to S8) to finally analyze the results.

**Figure S2.**
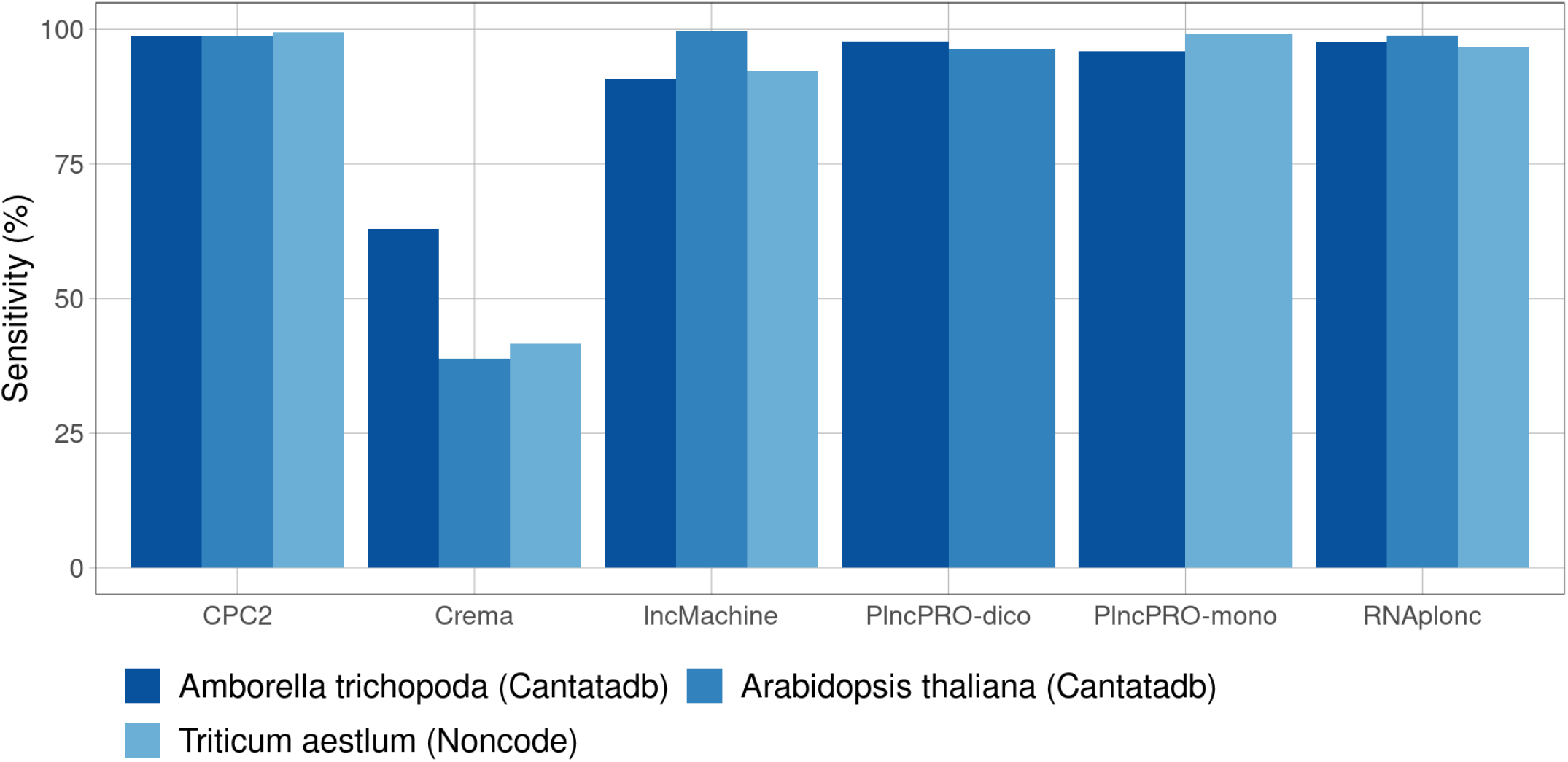
Performances of each tool applied on CANTATAdb and NONCODE datasets. Sensitivity values obtained by tools CPC2, Crema, lncMachine, PLncPro and RNAplonc on four plant datasets. The highest Sensitivity value (99.66%) was obtained with the lncMachine tool on dataset Arabidopsis thaliana (dicotyledonous). For the other three datasets, the best Sensitivity value (99.40%) was obtained with the CPC2 tool. Detailed values are available in Additional file 1: Table S6.

**Figure S3.**
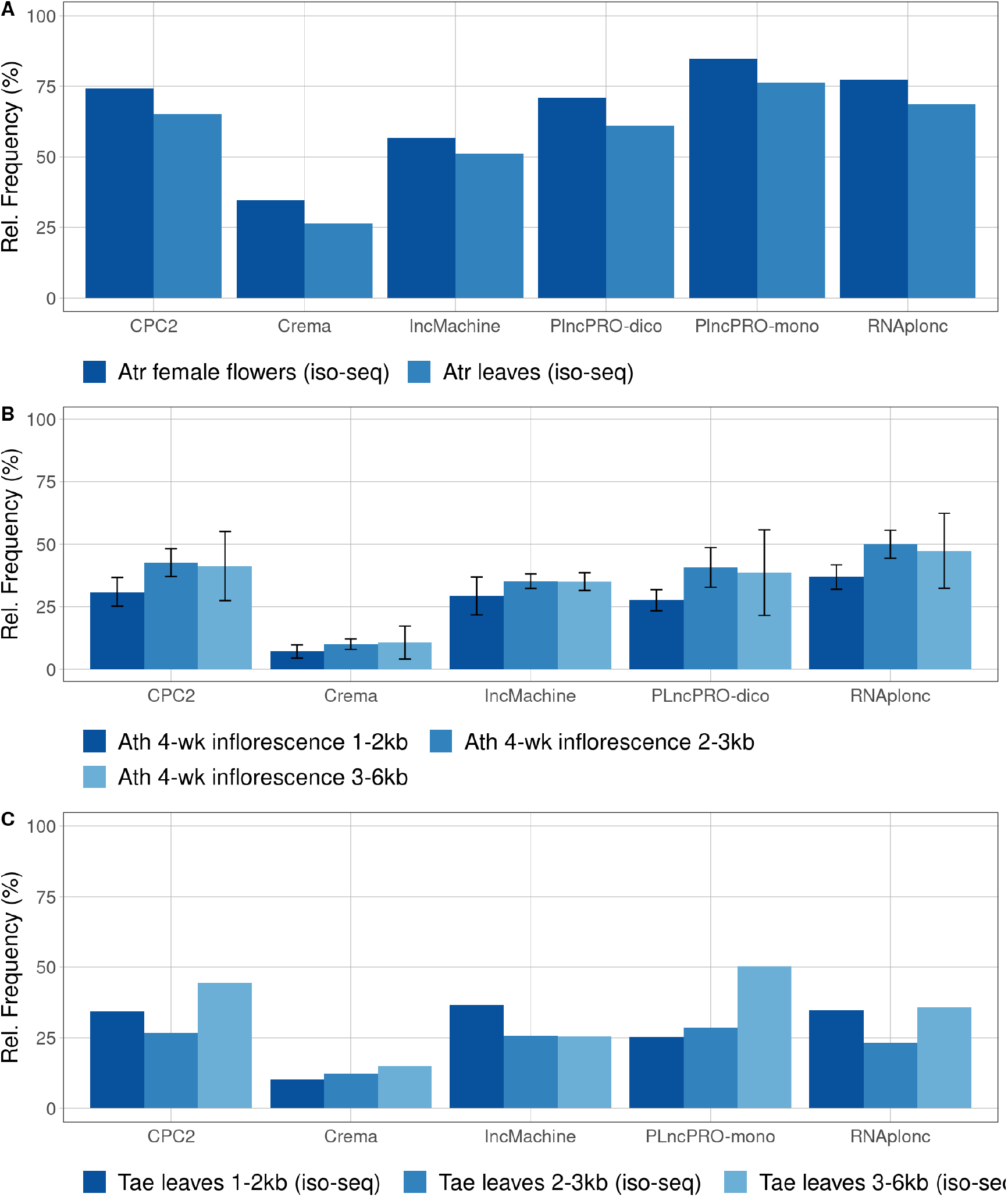
Relative frequency of lncRNAs on ISOdb plants datasets [23]. The 11 plant datasets from ISOdb were also analyzed using the five short-read data tools. Detailed values are available in Additional file 1: Table S7. A) Results for the datasets of Amborella trichopoda specie. The highest values of relative frequency were with the PLncPro (84.80%) and RNAplonc (77.18%) tools, for the monocotyledonous and dicotyledonous datasets, respectively. B) Results for the datasets of Arabidopsis thaliana specie. The tool with the highest relative frequency values was RNAplonc (57.97%), while for all tools the best values were obtained with the Ath 4-wk inflorescence 3-6kb rep2 dataset. C) Results for Triticum aestivum specie. The best result (36.64%) for the Tae leaves 1-2kb (iso-seq) dataset is for the lncMachine tool, while for the other two datasets it was obtained with the PLncPro tool.

**Figure S4.**
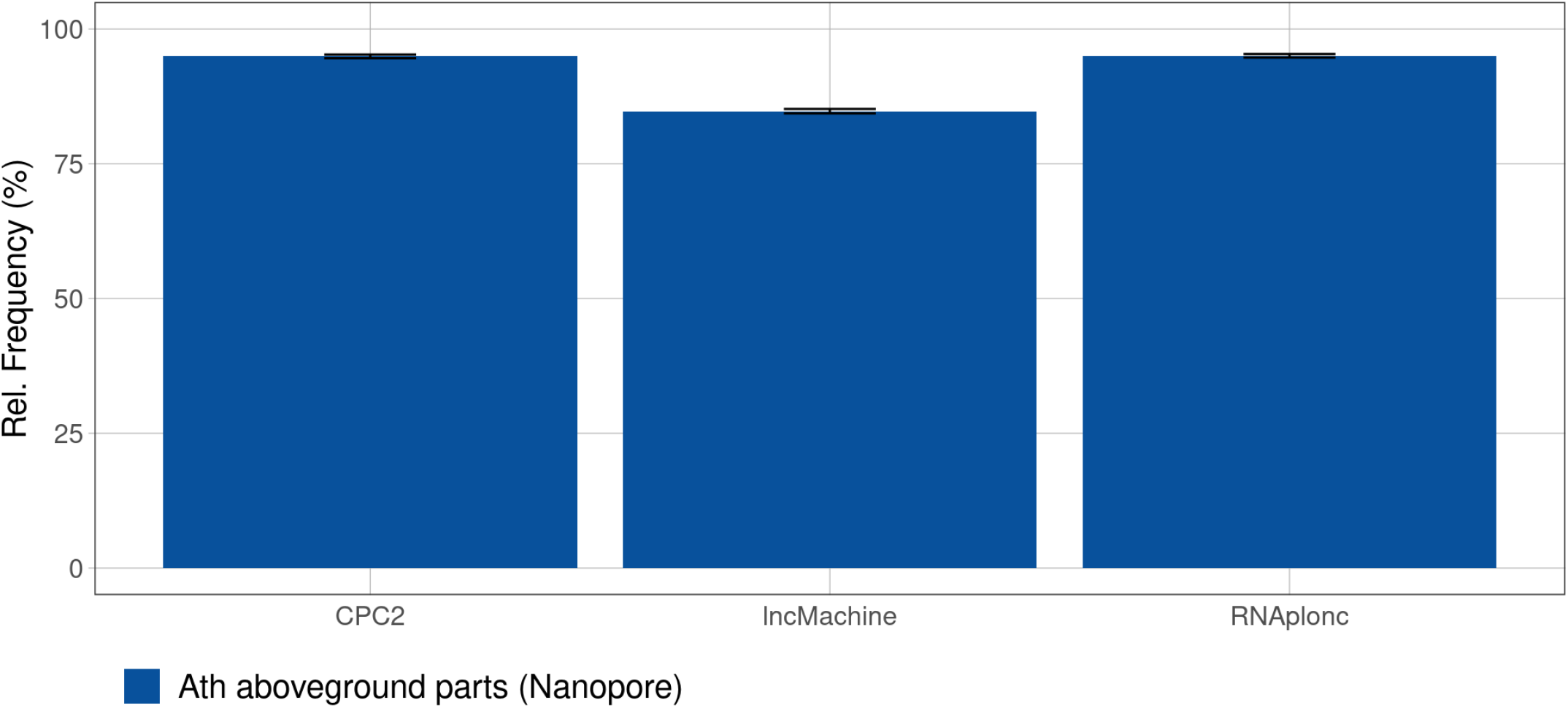
Relative frequency of lncRNAs on plants NANOPORE datasets. Relative Frequency obtained by CPC2, lncMachine and RNAplonc tools. The highest relative frequency value was obtained with the RNAplonc tool on all three plants NANOPORE datasets. Detailed values are available in Additional file 1: Table S8.

**Figure S5.**
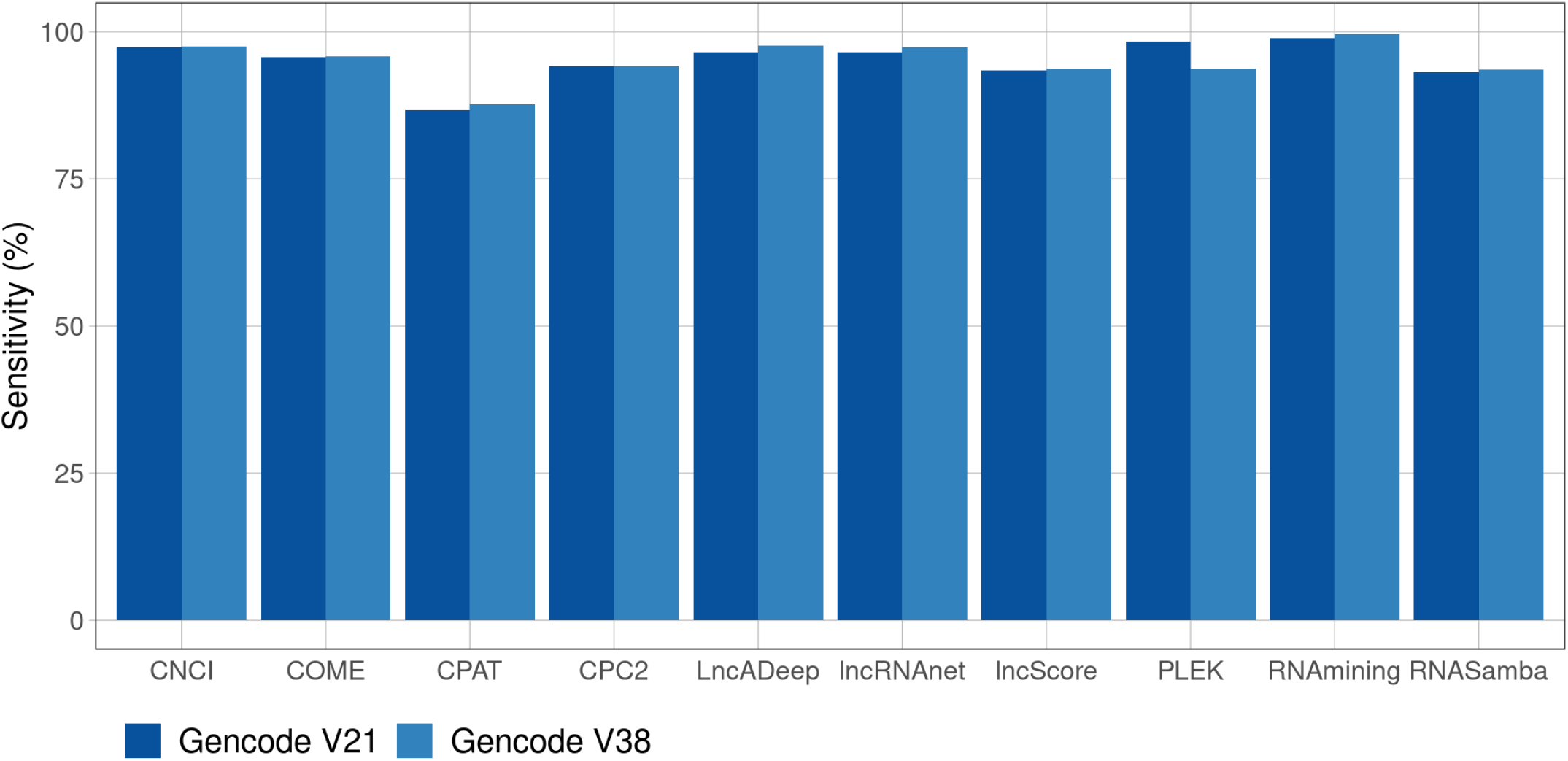
Performances of each tool applied on GENCODE v21 and v38 datasets. Sensitivity values obtained by tools CNCI, COME, CPAT, CPC2, LncADeep, lncRNAnet, lncScore, PLEK, RNAmining and RNASamba on GENCODE datasets. The highest Sensitivity value was obtained with the RNAmining tool on both datasets (98.88% and 99.56%). Detailed values are available in Additional file 1: Table S3.

**Figure S6.**
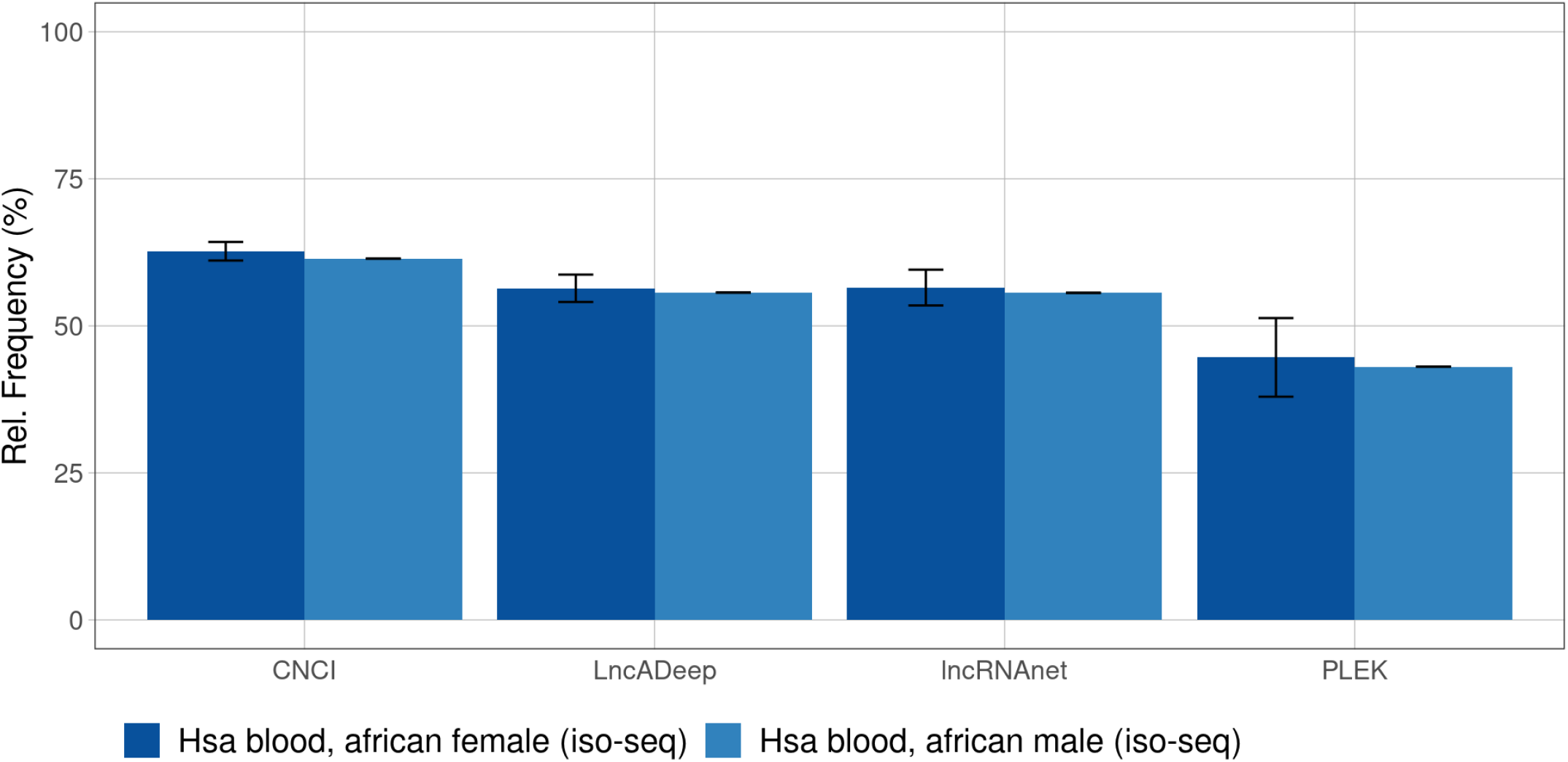
Relative frequency of lncRNAs on *Homo sapiens* on ISOdb Datasets. Relative Frequency obtained by CNCI, LncADeep, lncRNAnet and PLEK tools.The highest Relative Frequency value was obtained with CNCI on all two *Homo sapiens* ISOdb datasets (62.68% and 61.43%). Detailed values are available in Additional file 1: Table S4.

**Figure S7.**
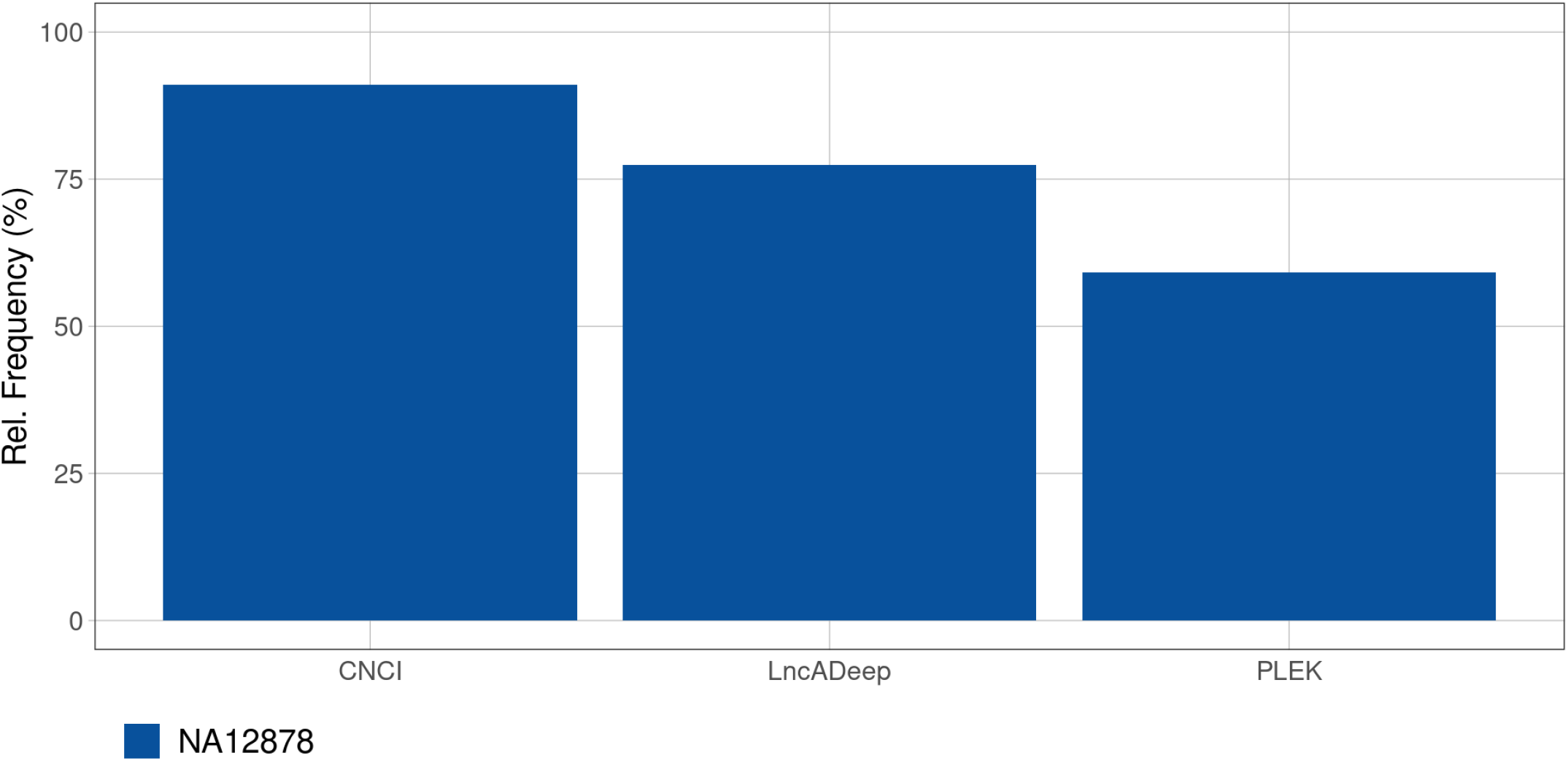
Relative Frequency of lncRNAs on *Homo sapiens* NANOPORE Dataset. Relative Frequency values obtained by CNCI, LncADeep and PLEK tools on Hsa reference sample dataset. The highest relative frequency value was obtained with the CNCI tool. Detailed values are available in Additional file 1: Table S5.

**Figure S8.**
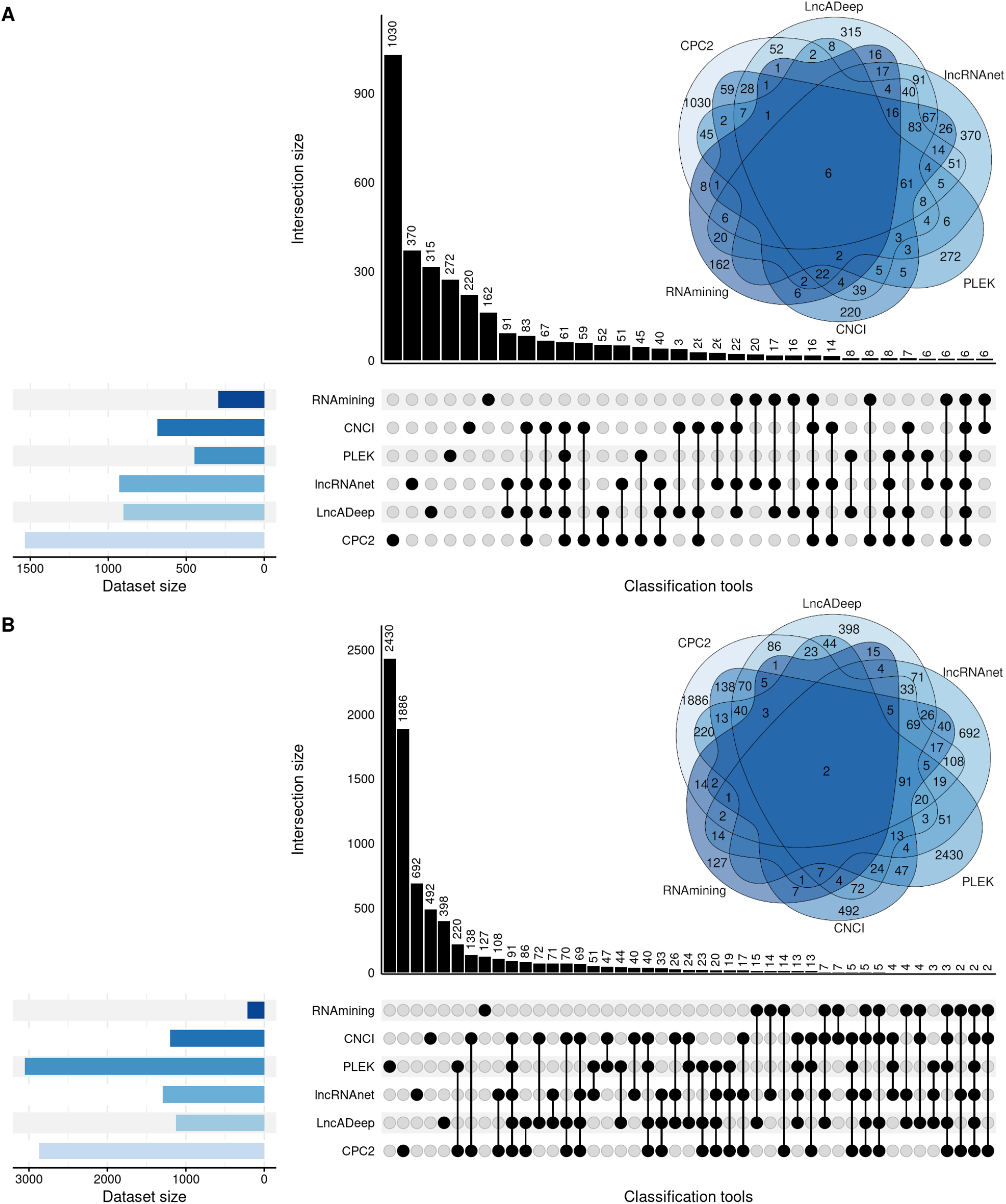
Upset plots for sequences classified as non-lncRNA by the 6 tools with the best average sensitivity. A) Six sequences were identified as non-lncRNA by all six tools in GENCODE version 21 dataset. B) Only two sequences were identified as non-lncRN by all 6 tools in GENCODE version 38 dataset.

