## Supplementary Material - Tables 1-8 for "Impact of sequencing technologies on long non-coding RNA computational identification"

### 1 Supplementary Tables

**Table 1.** The lncRNA tools overview used in this report for humans and plants

| Tools † | Version | Species | Programming language |
|---|---|---|---|
| CNCI[1] | 2 | *Homo sapiens* | Python |
| COME[2] | 11/2019 | *Homo sapiens* | R, Perl |
| CPAT[3] | 3.0.0 | *Homo sapiens* | Python, R |
| CPC2[4] | 1.0.1 | *Homo sapiens* / Plants | Python |
| CREMA[5] | 06/2020 | Plants | Python |
| LncADeep[6] | 1.0 | *Homo sapiens* | Python, R |
| LncMachine[7] | 0.1 | Plants | Python |
| LncRNAnet[8] | 08/2018 | *Homo sapiens* | Python |
| lncScore[9] | 1.0.2 | *Homo sapiens* | Python, Perl |
| PLEK[10] | 1.2 | *Homo sapiens* | Python |
| PLncPRO[11] | 1.2.2 | Plants | Python |
| RNAmining[12] | 1.0.4 | *Homo sapiens* | Python |
| RNAplonc[13] | 1.1 | Plants | Perl, Python |
| RNAsamba[14] | 0.2.5 | *Homo sapiens* | Python |

† Based on[15] categorization, the LncRNAnet has hybrid approach and the others Tools have a biological approach.

**Table 2.** The sequencing technologies datasets used for the evaluation of the lncRNAs tools

| Species | Sample Description | Ref | Sequences | Database | Data Type | HTS reads technology |
|---|---|---|---|---|---|---|
| *A. trichopoda* | Atr leaves | [16] | 91,576 | ISOdb | Transcripts | long |
| *A. trichopoda* | Atr female flowers | [16] | 55,723 | ISOdb | Transcripts | long |
| *A. trichopoda* | CANTATAdb 2.0 | [17] | 5,511 | CANTATAdb | lncRNAs | short |
| *A. thaliana* | Ath 4-wk inflorescence 1-2kb | [16] | 44,164 | ISOdb | Transcripts | long |
| *A. thaliana* | Ath 4-wk inflorescence 2-3kb | [16] | 54,648 | ISOdb | Transcripts | long |
| *A. thaliana* | Ath 4-wk inflorescence 3-6kb | [16] | 43,426 | ISOdb | Transcripts | long |
| *A. thaliana* | CANTATAdb 2.0 | [17] | 4,373 | CANTATAdb | lncRNAs | short |
| *A. thaliana* | Ath aboveground parts | [18] | 23,134,068 | SRA/NCBI | Transcripts | long |
| *H. sapiens* | Release 21 | [19] | 26,414 | GENCODE | lncRNAs | short |
| *H. sapiens* | Release 38 | [19] | 48,751 | GENCODE | lncRNAs | long |
| *H. sapiens* | Hsa reference sample | [20] | 10,302,647 | NANOPORE | Transcripts | long |
| *H. sapiens* | Hsa blood, african male | [16] | 33,651 | ISOdb | Transcripts | long |
| *H. sapiens* | Hsa blood, african female | [16] | 144,287 | ISOdb | Transcripts | long |
| *T. aestivum* | Tae leaves 1-2kb | [16] | 12,836 | ISOdb | Transcripts | long |
| *T. aestivum* | Tae leaves 2-3kb | [16] | 18,830 | ISOdb | Transcripts | long |
| *T. aestivum* | Tae leaves 3-6kb | [16] | 2,753 | ISOdb | Transcripts | long |
| *T. aestivum* | NONCODEv6 | [21] | 12,427 | NONCODE | lncRNAs | short |

**Table 3.** Performances of Each Tool Applied on GENCODE v21 and v38 Datasets

| Tools | v21 Short reads | | v38 Long reads | | Average | Gain/Lost |
|---|---|---|---|---|---|---|
| | Total | Sensitivity | Total | Sensitivity | | |
| CNCI | 26,413 | 97.40% | 48,751 | 97.54% | 97.47% | +0.14% |
| COME | 26,414 | 95.67% | 48,751 | 95.88% | 95.77% | +0.21% |
| CPAT | 26,414 | 86.74% | 48,751 | 87.67% | 87.20% | +0.93% |
| CPC2 | 26,414 | 94.19% | 48,751 | 94.11% | 94.15% | -0.08% |
| LncADeep | 26,413 | 96.58% | 48,751 | 97.68% | 96.96% | +1.10% |
| lncRNAnet | 26,412 | 96.48% | 48,749 | 97.34% | 96.91% | +0.86% |
| lncScore | 26,124 | 93.40% | 48,463 | 93.68% | 93.54% | +0.28% |
| PLEK | 26,414 | 98.28% | 48,470 | 93.70% | 95.99% | -4.58% |
| RNAmining | 26,414 | **98.88%** | 48,751 | **99.56%** | **99.22%** | +0.68 |
| RNASamba | 26,413 | 93.20% | 48,751 | 93.58% | 93.39% | +0.38% |
| Mean | | 92.47% | | 94.56% | 93.50% | +2.09% |

**Table 4.** Relative Frequency of lncRNAs on *Homo sapiens* on ISOdb Datasets

| Dataset | Sample | Total | CNCI | lncADeep | lncRNAnet | PLEK | RNAmining |
|---|---|---|---|---|---|---|---|
| ISOdb | SAMN00001694 | 64,638 | 61.57% | 54.74% | 54.36% | 39.91% | **97.92%** |
| ISOdb | SAMN00001695 | 33,651 | 63.80% | 58.03% | 58.66% | 49.36% | **97.58%** |
| ISOdb | SAMN00001696 | 79,649 | 61.43% | 55.66% | 55.62% | 43.05% | **97.71%** |

**Table 5.** Relative Frequency of lncRNAs on *Homo sapiens* NANOPORE Dataset

| Dataset | Total | CNCI | lncADeep | PLEK | RNAmining |
|---|---|---|---|---|---|
| NA12878 | 10,302,647 | 91.06% | 77.45% | 59.15% | **96.82%** |

**Table 6.** Performances of Each Tool Applied on CANTATAdb and NONCODE datasets

| Dataset | Total | RNAplonc | PLncPro-mono | PLncPro-dico | Crema | lncMachine | CPC2 |
|---|---|---|---|---|---|---|---|
| *Amborella trichopoda* | 5,511 | 97.59% | 97.71% | 95.84% | 62.87% | 90.73% | **98.66%** |
| *Arabidopsis thaliana* | 4,373 | 98.87% | - | 96.34% | 38.78% | **99.66%** | 98.61% |
| *Triticum aestivum* | 12,427 | 96.71% | 99.07% | - | 41.55% | 92.16% | **99.40%** |

**Table 7.** Relative Frequency of lncRNAs on ISOdb Plants Datasets[16]

| Dataset/Bio Sample | Total | RNAplonc | PLncPro-mono | PLncPro-dico | Crema | lncMachine | CPC2 |
|---|---|---|---|---|---|---|---|
| Amborella trichopoda | | | | | | | |
| SAMN06320627 | 91,576 | 68.58% | **76.32%** | 61.04% | 26.51% | 51.08% | 65.14% |
| SAMN06320628 | 55,723 | 77.18% | **84.80%** | 70.95% | 34.65% | 56.75% | 74.31% |
| Arabidopsis thaliana | | | | | | | |
| SAMN04456597 | 23,579 | **33.40%** | - | 24.55% | 5.24% | 23.97% | 26.88% |
| SAMN04456598 | 20,464 | **46.05%** | - | 35.11% | 8.55% | 33.16% | 38.69% |
| SAMN04456599 | 17,114 | **36.75%** | - | 26.54% | 6.04% | 32.56% | 31.46% |
| SAMN04456600 | 27,050 | **40.34%** | - | 30.57% | 8.94% | 34.62% | 34.99% |
| SAMN04456601 | 31,069 | **53.94%** | - | 46.33% | 11.47% | 37.31% | 46.56% |
| SAMN04456602 | 22,962 | **57.97%** | - | 50.76% | 15.33% | 37.56% | 50.99% |
| Triticum aestivum | | | | | | | |
| SAMN04456603 | 12,836 | 34.76% | 25.23% | - | 10.27% | **36.64%** | 34.54% |
| SAMN04456604 | 18,830 | 23.21% | **28.62%** | - | 12.36% | 25.68% | 26.76% |
| SAMN04456605 | 2,753 | 35.78% | **50.42%** | - | 15.03% | 25.50% | 44.57% |

**Table 8.** Relative Frequency of lncRNAs on Plants NANOPORE Datasets

| Dataset | Total | RNAplonc | lncMachine | CPC2 |
|---|---|---|---|---|
| SRR10611193 | 8,258,511 | **95.36%** | 85.19% | 95.24% |
| SRR10611194 | 7,849,574 | **94.69%** | 84.39% | 94.57% |
| SRR10611195 | 7,025,983 | **95.01%** | 84.69% | 94.93% |
